# Detecting regulatory elements in high-throughput reporter assays

**DOI:** 10.1101/2020.08.07.241901

**Authors:** Young-Sook Kim, Graham D. Johnson, Jungkyun Seo, Alejandro Barrera, William H. Majoros, Alejandro Ochoa, Andrew S. Allen, Timothy E. Reddy

## Abstract

High-throughput reporter assays such as self-transcribing active regulatory region sequencing (STARR-seq) have made it possible to measure genome-wide regulatory element activity across the human genome. The assays, however, also present substantial analytical challenges. Here, we identify technical biases that explain most of the variance in STARR-seq signals. We then develop a statistical model to correct those biases and to improve detection of regulatory elements. This approach substantially improves precision and recall over current methods, improves detection of both activating and repressive regulatory elements, and controls for false discoveries despite strong local signal correlations.

## Introduction

Gene regulation is of foundational importance to nearly all biological processes, and variation in gene regulatory activity plays a major role in human disease risk (Lee and Young 2013; Parker et al. 2013; Finucane et al. 2015). A major step toward measuring regulatory activity across the human genome has been the development of new high-throughput reporter assays such as STARR-seq (Arnold et al. 2013) that allow regulatory element activity to be quantified with high-throughput sequencing rather than with optical detection of a fluorescent or luminescent signal.

High-throughput reporter assays create substantial analytical challenges that are distinct from other genomic assays. There are significant local biases in high-throughput reporter assay signal that we show here are due to features of the underlying genomic sequence and experimental procedures. We show those local biases are robust across STARR-seq assays, suggesting that they are caused by the underlying genomic sequence influencing experiments or analysis. For example, nucleotide composition can alter PCR efficiency leading to under- and over-representation of some sequences. Meanwhile, highly repetitive sequences may not align uniquely to the human reference genome, thus biasing analyses.

A second analytical challenge is that, unlike for ChIP-seq analyses, STARR-seq signals can be both positive and negative, reflecting activation and repression, respectively. Meanwhile, unlike for RNA-seq analysis, the boundaries of regulatory elements are not pre-defined, and must be estimated from the data. Those challenges together impact signal representations and hinder robust estimation of regulatory activity from raw signals. The resulting distorted signals can lead to false positives and false negatives in the detection of regulatory elements.

To the best of our knowledge, there are currently no methods developed to identify and estimate the effect of both activating and repressing regulatory elements while correcting for the underlying sequence biases in high-throughput reporter assays. To overcome that challenge, our <u>c</u>orrecting reads and <u>a</u>nalysis of <u>d</u>ifferentially active <u>e</u>lements (CRADLE) model takes a two-step approach. First, CRADLE uses a mixture generalized linear regression model to estimate and correct major biases that we have identified in STARR-seq data. Next, CRADLE detects regions with statistically significant regulatory activity from the bias-corrected signals while rigorously controlling FDR. In doing so, CRADLE substantially improves the use of high-throughput reporter assay technologies by providing a robust estimation of regulatory activity and improved visualization of raw signals.

## Results

### DNA sequence biases STARR-seq signals

To identify sources of signal variance in STARR-seq, we analyzed data from two whole genome STARR-seq studies completed in different labs and in different human cell models: A549 (Johnson et al. 2018) and HeLa-S3 cells (Muerdter et al. 2018). Each study followed a similar protocol in which an input STARR-seq library was generated by cloning randomly fragmented genomic DNA into the 3’ untranslated region (UTR) of a reporter gene. The input library was then assayed by transfecting it into cultured human cells where the cloned DNA fragments regulate their own transcription into mRNA. The expression of each random fragment as mRNA was then measured with high-throughput sequencing. Finally, regulatory activity was estimated by comparing the expression of each fragment in the output library relative to its abundance in the input library.

The STARR-seq input libraries exhibited substantially more signal variance than is observed in the controls for other genomic assays such as for ChIP-seq input (Fig. 1A-B; (Consortium 2012)). That variance in STARR-seq input signal was consistent across replicates and between studies (Fig. 1C), and more so than for ChIP-seq input signal (Fig. S1). Here, we analyzed four potential sources of variance in STARR-seq signal: (i) DNA structure influencing DNA fragmentation, cloning or other enzymatic reactions, thus affecting which DNA fragments are available in the assay library (Poptsova et al. 2014); (ii) differences in the Gibbs free energy of DNA fragments influencing multiplex PCR efficiency, leading to preferential amplification of some fragments (Cheung et al. 2011; Benjamini and Speed 2012; Hansen et al. 2012; Jiang et al. 2015; Love et al. 2016; Teng and Irizarry 2017); (iii) G-quadraplexes in the genome impairing amplification by DNA polymerase (Chambers et al. 2015; Rhodes and Lipps 2015); and (iv) biases resulting from differences in the mappability of short read sequences to the reference human genome, for example due to repetitive sequences (Derrien et al. 2012).

**Figure 1.**
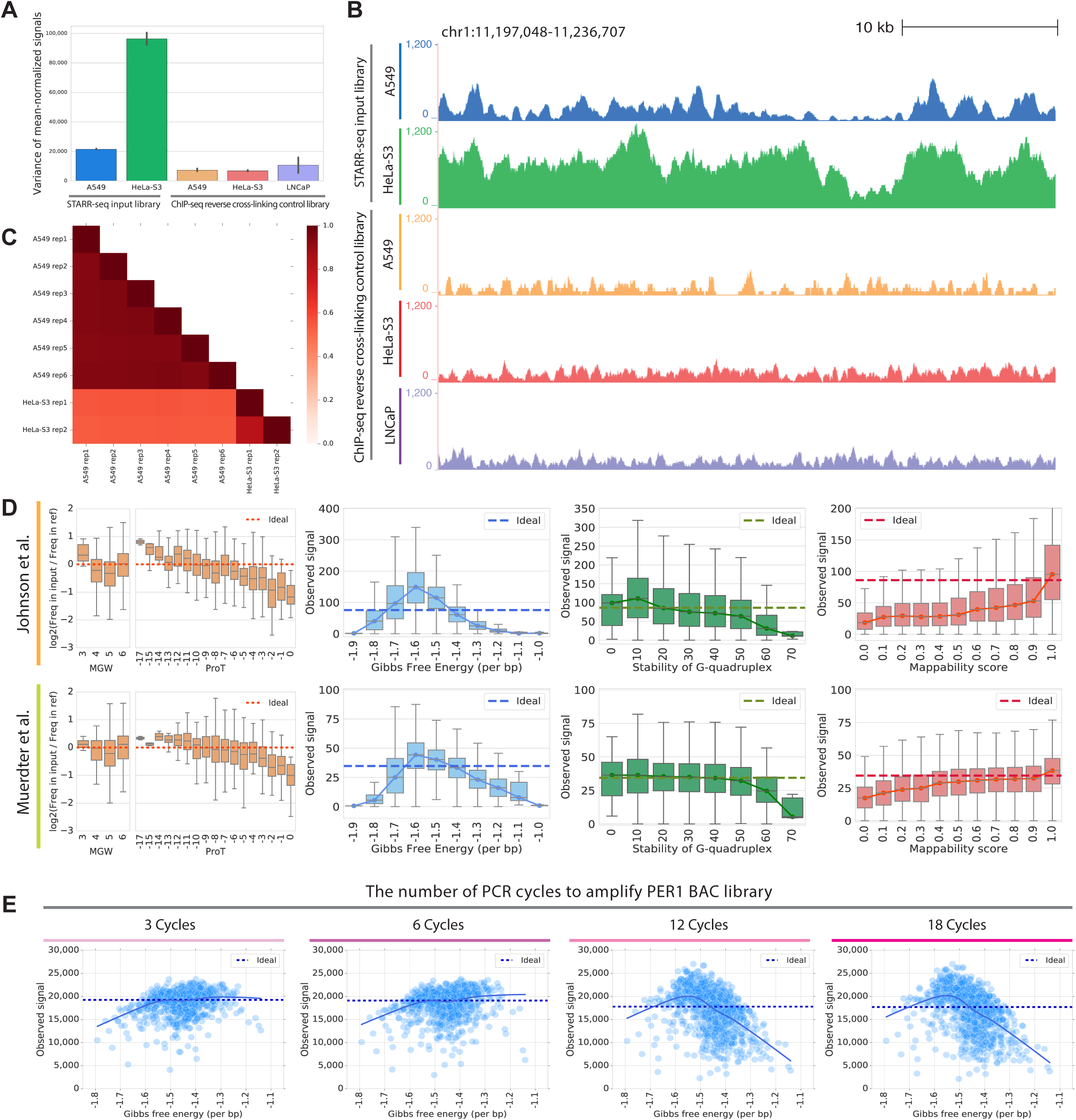
Technical biases affect STARR-seq signal. (A) Signal variance in STARR-seq input libraries exceeds that observed in ChIP-seq input control libraries. Variance in per base signal in individual RPKM-normalized libraries are plotted for chromosome 1. The error bars indicate variance between replicates. (B) Representative browser signal tracks for STARR-seq and ChIP-seq input libraries (chr1:11197048-11236707). Signals are RPKM-normalized. (C) Pearson correlations of STARR-seq input library signals (Johnson et al. 2018; Muerdter et al. 2018) in 1 bp windows along chromosome 1. (D) DNA sequence biases impact STARR-seq signals. STARR-seq signals are plotted for 500 bp windows with varying degrees of bias for the following physical properties of DNA: fragment-end DNA structures, Gibbs free energy, G-quadruplex structure, and mappability. In plots of fragment-end bias, the ideal line is y=0. In plots of other biases, the ideal line is the median signal. Whiskers extend 1.5 times the inter-quartile range. Center lines show the medians. MGW, minor groove width; ProT, propeller twist. (E) PCR amplification introduces bias into STARR-seq libraries. Impact of Gibbs free energy bias on PER1 BAC libraries amplified with different number of PCR cycles (3, 6, 12, and 18 cycles). Each point represents the sum of signals in a 500 bp window from three technical replicates. The solid line is a lowess fit line. The dashed ideal line is the median signal across all windows.

We found evidence that each source of bias influences the signal observed when sequencing STARR-seq libraries. To model biases due to DNA secondary structure, we computationally estimated the minor groove width (MGW) and propeller twist (ProT) at the 5’ ends of DNA fragments (Zhou et al. 2013). We analyzed 5’ ends of DNA fragments because they are not modified in the end-repairing process of generating STARR-seq libraries (Poptsova et al. 2014). We observed distinct biases in 5’ MGW and ProT (Fig. 1D). Consistent with preferential fragmentation partially contributing to that bias, we found that ApG and GpG dinucleotides are most underrepresented at the 5’ ends of STARR-seq fragments, which were previously reported to be less prone to shearing (Fig. S2; (Poptsova et al. 2014)). To estimate biases due to changes in Gibbs free energy, G-quadruplex structure, and mappability, we binned the genome into 500bp windows and used empirical datasets to estimate the Gibbs free energy (Protozanova et al. 2004), stability of G-quadruplex structure (Chambers et al. 2015), and the fraction of redundant mappable positions in the reference genome for each window (Derrien et al. 2012), respectively (Fig. 1D). Fragments with extreme Gibbs free energy, highly stable G-quadruplex structure, and low mappability all had substantially depleted STARR-seq signals. Those trends were generally consistent across two whole genome STARR-seq studies (Johnson et al. 2018; Muerdter et al. 2018).

To confirm that biases in estimated Gibbs free energy are due to differences in PCR amplification efficiency, we generated DNA fragment libraries from a bacterial artificial chromosome (BAC) using between three and 18 cycles of PCR. The BAC contained 211 kb from human chromosome 17. We observed the same trend in that DNA fragments with extreme Gibbs free energy were depleted from the final library (Fig. 1E). Further, the magnitude of that bias increased substantially with additional cycles of PCR, particularly for fragments with the highest Gibbs free energy. That trend also informs the need of independently estimating biases in each library because output STARR-seq libraries are likely to have more severe PCR-related biases than input libraries due to additional PCR cycles required (Johnson et al. 2018; Muerdter et al. 2018).

### Modeling biases in STARR-seq signal guides improved experimental designs

To model the above biases in STARR-seq signal, we developed a generalized linear regression model (GLM) with covariates to model DNA structure (Zhou et al. 2013) in fragment-ends, annealing and denaturing efficiency of DNA fragments related to their Gibbs free energy (Protozanova et al. 2004), stability of G-quadruplex structure (Chambers et al. 2015), and mappability (Derrien et al. 2012) as a reduced set of independent variables (Fig. 2A). We then fit that model to predict biases in STARR-seq signals across the genome (Fig. 2A). Here, we took two approaches to improve model fit: 1) a mixture model and 2) a structured sampling of a training set. The use of a mixture model was motivated by our observations that regions with high STARR-seq signal have significantly different coefficients from the rest of regions, especially for independent variables related to Gibbs free energy bias (Fig. 2B). This led us to use a mixture model that estimates bias impact separately for regions with high signal. To better fit the tails of the distribution, we used a structured sampling approach that selected a pre-defined percentage of a training set in a given quantile range of signal (Fig. 2C). Together, these adjustments better modeled the tails of the STARR-seq signal distribution where biases are most strong and thus most likely to impact analysis.

**Figure 2.**
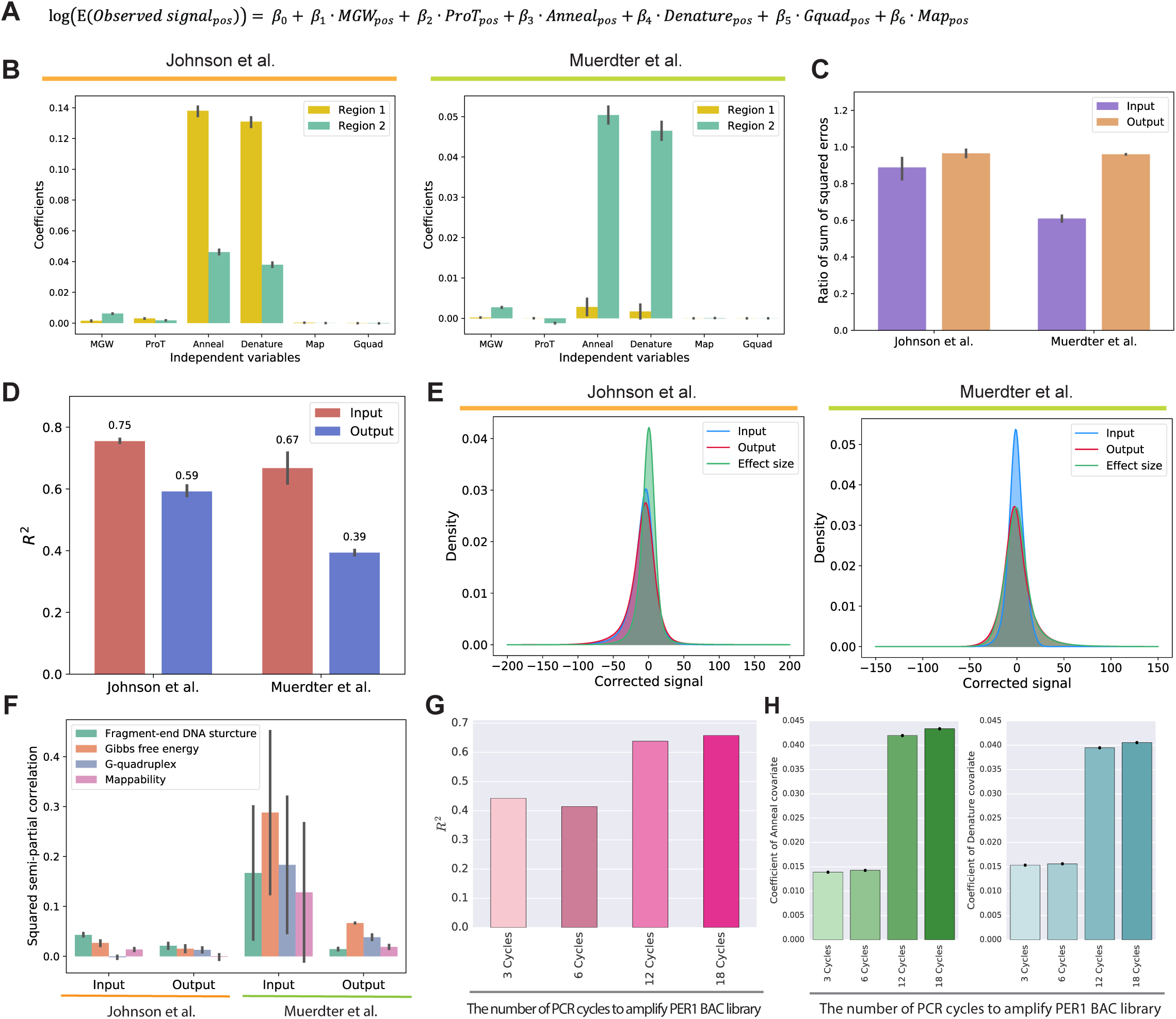
The CRADLE GLM approach accurately predicts signal bias. (A) Equation of the GLM to predict the impact of technical biases. Notation: pos, single-bp position; MGWpos, minor grove width; ProT_pos_, propeller twist; Anneal_pos_, annealing efficiency; Denature_pos_, denaturation efficiency; Gquad_pos_, G-quadruplex structure; Map_pos_, mappability. (B)-(F) The results from the GLM fitted with the STARR-seq input libraries and the 0hr-dex-treated (Johnson et al. 2018) and no-inhibitor-treated output libraries (Muerdter et al. 2018) are visualized for chromosome 1. (B) Coefficients in input libraries for regions with signals below and above 90th percentile (‘Region 1’ and ‘Region 2’, respectively). (C) Ratio of sum of squared errors with structured sampling to sum of squared errors with random sampling for regions with extremely high signals (above 99th percentile). (D) Variance explained by CRADLE. The R2 values from GLMs fitted with input and output STARR-seq libraries. The error bars indicate variance between replicates. (E) Distribution of GLM residuals and the STARR-seq effect sizes after correction. (F) Squared semi-partial correlations of fragment-end, Gibbs free energy, G-quadruplex, and mappability covariates. The error bars indicate variance between replicates. (G) The R2 of the GLM fitted with PER1 BAC libraries amplified with different number of PCR cycles. (H) Coefficients of anneal and denature covariates from the GLM fitted with PER1 BAC libraries. The error bars show 95% confidence interval.

Overall, the GLM fit the observed signals with R^2^ up to 0.75 for input STARR-seq libraries (Fig. 2D). The model fit output libraries less well, ostensibly due in part to regulatory activity also contributing to differences in STARR-seq signal. Residuals from the model approximately follow a normal distribution (Fig. 2E), supporting model fit. We also estimated the extent to which each covariate uniquely explained variation in STARR-seq signal (Fig. 2F). Overall, the median of the explained variation across the two studies showed fragment-end and Gibbs free energy bias explained the greatest amount of signal variation. In the Johnson et al. dataset (Johnson et al. 2018), G-quadruplex bias in the input STARR-seq library and mappability bias in the output STARR-seq library had a negative marginal contribution to total predictive power but the effects were minor. Meanwhile, the full model explains most of the signal variation in the STARR-seq data.

Most of the parameters we modeled are not readily mitigated by modifying experimental procedures. For example, reducing PCR cycles to mitigate PCR biases may not be feasible when template is limited. Therefore, we investigated whether the mixture GLM can instead be used to statistically correct biases in STARR-seq signal. First, we fit the above mixture GLM to fragment sequencing libraries generated with different numbers of PCR cycles, and calculated the amount of variance explained by the mixture GLM. Consistent with our earlier observation that additional PCR cycles increased Gibbs free energy bias (Fig. 1E), the model explained more signal variance when more PCR cycles were used (Fig. 2G). There was also a monotonic increase in the coefficients for fragment annealing and denaturing efficiency based on the Gibbs free energy (Fig. 2H). Those results of bias predictions demonstrate that the mixture GLM can correct different amounts of bias related to different experimental designs.

### Removing technical biases in STARR-seq improves visualization

Because most signal variation in STARR-seq is due to the underlying DNA sequence, it is challenging to visually inspect uncorrected STARR-seq signals and assess the quality of experiments and of statistical analyses. That visualization can be substantially improved by instead using the residuals from the mixture GLM model where those sources of local signal variation have been removed. For example, across the two genome-wide studies analyzed here (Johnson et al. 2018; Muerdter et al. 2018), the mixture GLM reduced signal variance by between 40 and 80% with approximately zero-centered corrected signals (Fig. 3A-B). Further demonstrating generality across specific experimental procedures, the mixture GLM also effectively corrected biases due to different amounts of PCR (Fig. 3C-D). An example of the resulting correction for a 19 kb region is demonstrated in Fig. 3E, where the GLM residuals allows for clearer visual identification of elements with activating or repressive regulatory effects. For example, a region near the PER1 gene was previously identified as having activating regulatory activity in response to dexamethasone (dex) treatment that binds to glucocorticoid receptor (GR) (Reddy et al. 2012; Johnson et al. 2018). This region showed much clearer indication of activity after correction (Fig. 3F). Similarly, an example of a repressive element that is bound by the REST repressor (Consortium 2012) is also better represented in corrected signals compared to uncorrected observed signals (Fig. 3G). Together, these results demonstrate that our model accounts for a substantial variation of signals in STARR-seq data and improves visualization of signals.

**Figure 3.**
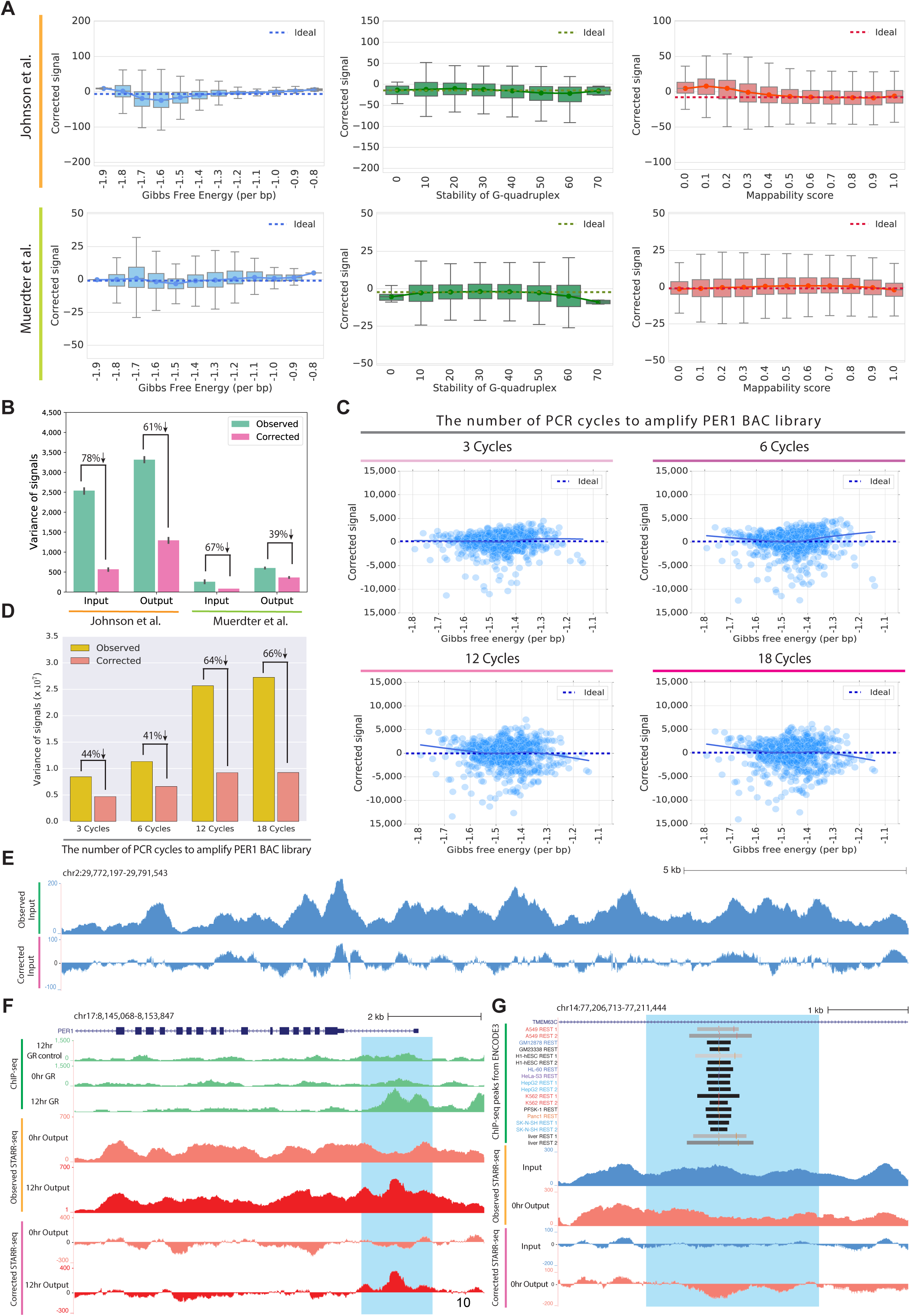
The CRADLE GLM approach corrects technical bias. (A) STARR-seq signals are plotted for 500 bp windows along chromosome 1 after removing technical bias with CRADLE. Signal is balanced despite varying degrees of technical biases. The ideal line is the median corrected signal. Whiskers extend 1.5 times the interquartile range. Center lines show the medians. (B) Variance in observed signals and CRADLE corrected signals in 1 bp windows along chromosome 1. The error bars indicate variance between replicates. (C) STARR-seq signals in PER1 BAC libraries amplified with different number of PCR cycles after removing technical bias with CRADLE. Each point represents the sum of corrected signals in a 500 bp window from three technical replicates. The solid line is a lowess fit line. The dashed ideal line is the median signal across all windows. (D) Variance in observed signals and CRADLE corrected signals in 1 bp windows after correcting the PER1 BAC libraries. The error bars indicate variance between replicates. (E) Representative signal tracks of ST ARR-seq input libraries before and after CRADLE correction (chr2:29779750-29786199). (F) STARR-seq and ChIP-seq signal tracks in the dex-responsive PER1 locus. Observed and corrected STARR-seq signals are presented for 0 hr dex-treated (untreated) and 12 hr dex-treated output libraries (Johnson et al. 2018). ChIP-seq signal tracks are not corrected. The highlighted region (chr17:8151204-8152809) is a known dex-responsive activating regulatory element. (G) STARR-seq signal tracks in the TMEM63C locus. Observed and corrected STARR-seq signals are presented for input and 0 hr dex-treated output libraries (Johnson et al. 2018). The highlighted region (chr14:77207895-77210261) contains a REST motif and is occupied by REST in multiple cell types.

### Correcting biases improves detection of regulatory signals embedded in STARR-seq data

STARR-seq measures both activation and repression of reporter gene expression (Johnson et al. 2018). However, statistical models for ChIP-seq or ATAC-seq are typically designed to detect only increases in signal. At the same time, local STARR-seq signals are highly correlated (e.g. Fig. 3E). That correlation, if not appropriately considered, can lead to nonconservative control of type I errors (Lun and Smyth 2014). To overcome those challenges, we developed a two-step statistical approach to merge locally correlated signals while maintaining FDR control (Fig. 4A; see Methods). Briefly, our approach first detects signals in broad genomic regions, and then identifies more specific sources of signal variation within those regions. The approach is based on previous work from Benjamini (Benjamini and Hochberg 1995; Benjamini and Bogomolov 2014). To increase power of detecting regulatory elements in this approach, we applied independent filtering to remove regions without enough signal variation to reject the null (Fig. 4A-B; (Bourgon et al. 2010)).

**Figure 4.**
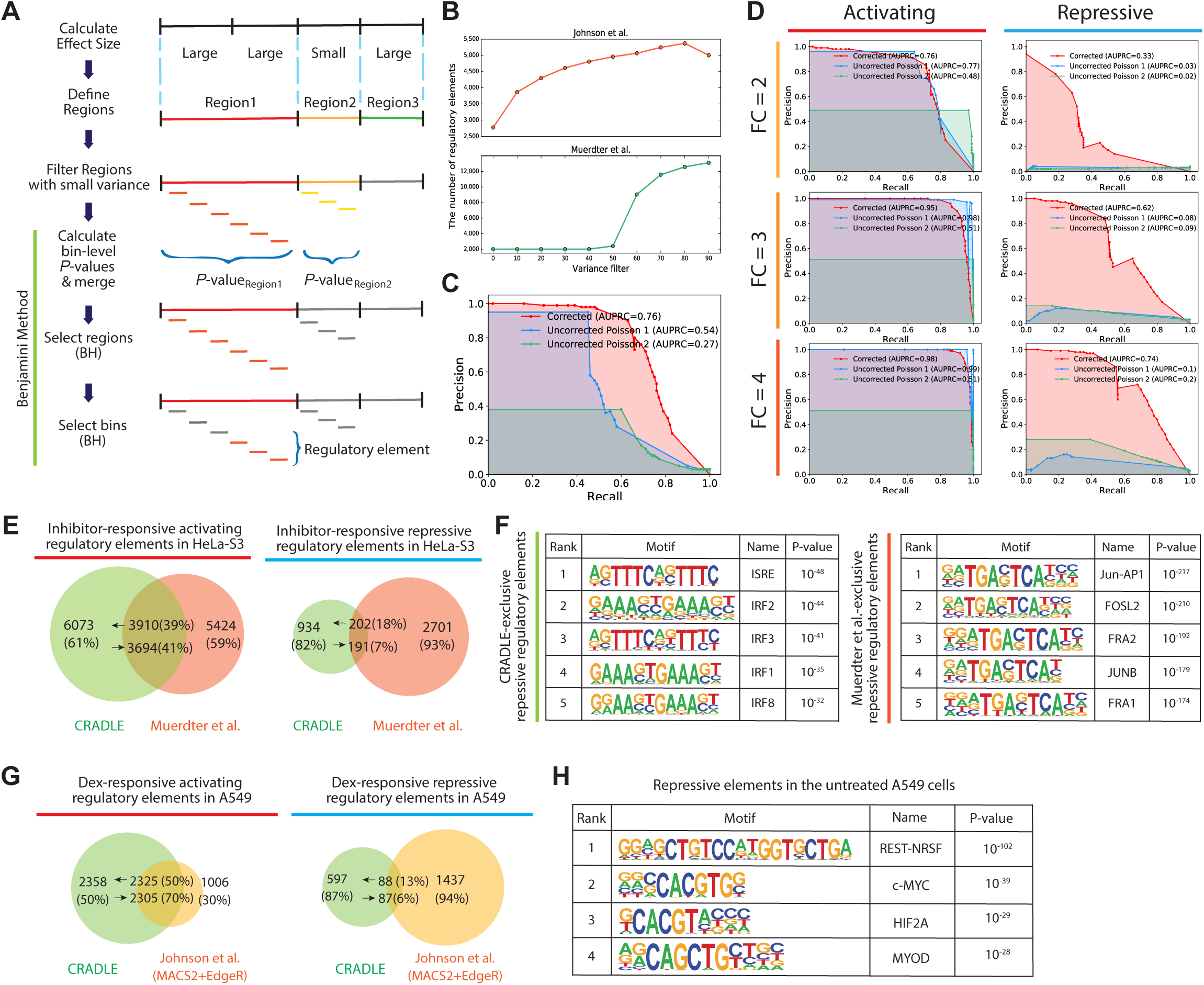
Detection of regulatory elements with CRADLE. (A) CRADLE regulatory element pipeline. Effect sizes are calculated in windows of uniform length. Contiguous windows with similar effect sizes are merged into regions prior to filtering regions with small variance. Regions are binned and a statistical test is performed on each bin to compare corrected input and output signals. Bin-level P-values are merged to generate a region-level P-value prior to performing a region-level Benjamini-Hochberg (BH) procedure. Regions selected by the first BH procedure were used to perform a bin-level BH procedure to identify regulatory elements. (B) The number of detected regulatory elements is dependent on the variance filter. (C)-(D) Precision-recall curves, using corrected and uncorrected signals in the simulation study. To detect regulatory elements with uncorrected signals, two statistical approaches were used: 1) fitting uncorrected signals to Poisson GLM and performing Wald test (‘Uncorrected Poisson 1’) and 2) Using a Poisson distribution with the mean of uncorrected input signals as a null distribution and testing the significance of the mean of uncorrected output signals (‘Uncorrected Poisson 2’). (C) Precision-recall curve when signals are simulated with mixed fold change (2, 3, 4) and a mix of activating and repressive elements. (D) Precision-recall curve when signals are simulated with a fixed fold change (FC) and with a fixed regulatory activity (either activating or repressive). (E)-(F) Comparison of inhibitor-responsive regulatory elements detected by CRADLE and Muerdter et al. (Muerdter et al. 2018) (E) The overlap of regulatory elements detected by both studies. (F) Transcription factor motif enrichment in repressive regulatory elements exclusively detected by each study. (G) The overlap of dex-responsive activating and repressive regulatory elements detected by CRADLE and Johnson et al. (Johnson et al. 2018). (H) Transcription factor motif enrichment in A549 steady-state repressive regulatory elements detected by CRADLE.

To demonstrate the benefit of correcting technical biases when detecting regulatory elements in STARR-seq data, we simulated whole genome STARR-seq signals with embedded activating and repressive regulatory elements across a range of effect sizes (Fig. 4C-D). We then used the method described above to detect regulatory elements in corrected or uncorrected signals. When detecting regulatory elements with uncorrected signals, we changed the statistical testing in the method described above from t-test that requires normality assumption to two statistical approaches that used a Poisson distribution to avoid unfairly reducing the performance with using uncorrected signals. Specifically, we used following two approaches: 1) fitting uncorrected signals to Poisson GLM and performing Wald test (‘Uncorrected Poisson 1’) and 2) Using a Poisson distribution with the mean of uncorrected input signals as a null distribution and testing the significance of the mean of uncorrected output signals.

Overall, correcting biases with the GLM substantially improved the precision of detecting regulatory signals, especially at more stringent detection thresholds (Fig. 4C-D). In contrast, with uncorrected signals, the majority of regulatory elements were false positives (Fig. S3). The improved performance was particularly pronounced for detecting repression (Fig. 4D), demonstrating that removing technical biases is particularly important for detecting repressive regulatory element activity. The increase of area under the precision recall curve (AUPRC), using corrected signals was up to 0.64, compared to using uncorrected signals.

Overall, repressive signals were more difficult to detect. In the repression simulation with corrected signals, recall and precision were significantly smaller than activation simulation by up to 0.43 in AUPRC. This decreased AUPRC was mainly due to small simulated output signals of repressive regulatory elements that were largely filtered out by the overall variance filter. However, this simulation result still shows correcting technical biases helps to decrease false positives in detecting both activation and repression.

### Improved detection of regulatory elements in STARR-seq data

We used the method described above to call regulatory elements in data from two published whole genome STARR-seq studies (Johnson et al. 2018; Muerdter et al. 2018). Muerdter et al. measured differential regulatory activity in response to inhibitors that blocked interferon response (Muerdter et al. 2018). The study reported 12,010 inhibitor-responsive regulatory elements with 2,892 repressive elements, with their analysis pipeline that used binomial distribution and hypergeometric tests (Arnold et al. 2013; Muerdter et al. 2018). CRADLE detected a similar number of regulatory elements at 20% FDR (N = 11,119), 1,136 of which were repressive elements (Table S1). While the activating elements detected by each method had substantial overlap, repressive elements were largely different between the methods (Fig. 4E).

To investigate the biological properties of regulatory elements detected exclusively by CRADLE and the method used in Muerdter et al. (Muerdter et al. 2018), we performed motif enrichment analysis on the non-overlapping regulatory elements. As expected from the experimental design, motifs for interferon-responsive transcription factors (TFs) were most strongly enriched in the CRADLE-exclusive repressive elements (Fig. 4F, Table S2). In contrast, AP-1 TF motifs were most significantly enriched in the repressive elements unique to the Muerdter et al. analysis (Muerdter et al. 2018), followed by interferon response motifs lower in the ranking (Fig. 4F, Table S3). We noted that CRADLE estimated positive effects for 1,692 repressive elements uniquely detected by Muerdter et al. (Fig. S4; (Muerdter et al. 2018)), suggesting they are false positives due to the biases in STARR-seq signal. Indeed, subsequent motif analysis of those repressive elements with positive effects revealed enriched NF-kB motifs, not corresponding to the experimental design.

The Johnson et al. study used STARR-seq to measure changes in regulatory activity in response to the dex across time (Johnson et al. 2018). The study used MACS2 (Zhang et al. 2008) and edgeR (Robinson et al. 2010) together to identify 4,835 dex-responsive regulatory elements at 0.05 FDR with 3,311 activating elements. With the data from Johnson et al. (Johnson et al. 2018), we used CRADLE to detect regulatory elements both in untreated A549 cells and in response to dex at the same 0.05 FDR (Table S4-S5). That analysis identified 10% more dex-responsive regulatory elements (N = 5,368) than the methods used by Johnson et al. (Johnson et al. 2018), with 4,683 activating and 685 repressive dex-responsive regulatory elements (Table S5). As with the comparison to the Muerdter et al. analysis, we observed little overlap in repressive elements while activating elements showed up to 70% overlap (Fig. 4G). Overall, those repressive regulatory elements identified by CRADLE in each study had higher control library signals than activating regulatory elements, demonstrating CRADLE requires a region to have enough coverage to be reliably detected as repressive (Fig. S5).

To validate the newly identified repressive elements, we performed motif enrichment analysis to determine enriched motifs for repressive elements. The motif for the RE1-silencing transcription factor (REST), a well-characterized repressive factor (Chong et al. 1995), was most enriched in repressive regulatory elements in untreated A549 cells, followed by TFs binding to the E-box motif (Fig. 4H, Table S6). Meanwhile, the motif enrichment in dex-responsive activating and repressive elements from CRADLE corresponded to previous findings about GR biology (Table S7-S8). Namely, for the dex-responsive activating regulatory elements, the GR DNA binding motif was most enriched followed by co-factor AP1 transcription family (Johnson et al. 2018) (Table S7). For the dex-responsive repressive regulatory elements, the AP1 motif was most enriched, consistent with the role of AP1 in GR-mediated activation and repression (Table S8; (Gupte et al. 2013; Johnson et al. 2018; McDowell et al. 2018)).

We also validated some of the 240 A549 steady-state REST-binding repressive regulatory elements using two independent studies (Fig. S6; (van Arensbergen et al. 2019; Doni Jayavelu et al. 2020)). Though neither of these studies used A549 cells, we assumed the repressive activity of the REST-binding repressive regulatory elements could be validated in other cell models because REST is a common repressor in diverse cell lines. We intersected the REST-binding repressive regulatory elements with the regions tested by Jayavelu et al. (Doni Jayavelu et al. 2020) that used massively parallel reporter assay (MPRA) test repressive activity, and observed 30 elements in common (Fig. S6). Of those, 27 (90%) elements had repressive activity in Jayavelu et al. while two elements did not have coverage and one element did not exhibit repressive activity. The one non-repressive element is likely due to the small overlap with their tested region (27bp) that did not cover the REST motif. We also compared our repressive element calls with the data from a genome-wide survey of regulatory elements (SuRE) signal (van Arensbergen et al. 2019). The SuRE signal in the REST-binding repressive regulatory elements exhibited repression in that study that was significantly different from both random and activator-binding elements (Fig. S6).

## Discussion

We demonstrated the majority of variation in STARR-seq signal can be explained by DNA-sequence features that are related to experimental biases rather than regulatory element activity. Overall, biases in PCR amplification had some of the strongest impacts on sequence biases in STARR-seq, and we show here that minimizing the amount of PCR can dramatically reduce variation in signals. DNA structure bias at the ends of fragments is possibly caused by preferential fragmentation, cloning, or efficiency as an enzymatic substrate. The efficiency of adding adaptors in cloning or in reverse transcription could be also affected by DNA sequences or structures at fragment-ends (Zheng et al. 2011). Potential opportunities to mitigate those biases could include using multiple enzymes from different species that have different sequence biases, or further refinement of reaction conditions. Similarly, increasing read length could mitigate mappability-induced biases by decreasing the mappable space in the genome; and G-quadruplex structure bias might be alleviated by optimizing experimental conditions to de-stabilize those structures.

However, mitigating technical biases by optimizing experiment conditions is not always feasible. That motivated us to develop a statistical model to computationally mitigate those biases. We demonstrated the mixture GLM had significant predictive power that led to substantially stabilized STARR-seq signals. Indeed, corrected signals showed noticeably reduced variance and improved visualization of regulatory activity.

With corrected signals from the mixture GLM, we detected regulatory elements with substantially improved accuracy compared to previous models. CRADLE especially improved the identification of repressive regulatory elements that were challenging to detect previously, as we demonstrated via simulations, comparisons to other studies, and through investigation of DNA binding motifs for repressive factors. That improvement will allow for a more complete understanding of the diversity of regulatory element activity across the human genome.

To the best of our knowledge, CRADLE is the first statistical model that corrects biases and detects regulatory elements in STARR-seq. We expect CRADLE serves as a powerful statistical tool to discover both activating and repressive regulatory elements in STARR-seq with increased sensitivity and precision. This model also provides an opportunity to re-assess preexisting STARR-seq data and discover novel regulatory elements. CRADLE is implemented in Python and freely available at https://github.com/Young-Sook/CRADLE.

## Methods

### Downloaded data

For STARR-seq data, we downloaded fastq files of whole genome STARR-seq that used A549 and HeLa-S3 cells from Johnson et al. (Johnson et al. 2018) and Muerdter et al. (Muerdter et al. 2018), respectively. Those files were downloaded from NCBI Gene Expression Omnibus (GEO) repository (Barrett et al. 2013) with accession codes available in those studies (GSE114063, GSE100432).

For ChIP-seq data (Consortium 2012; Davis et al. 2018), we downloaded ChIP-seq fastq files with following GEO accession codes: GSE91296 for A549 control ChIP-seq; GSE91275 for A549 0hr-dex-treated GR ChIP-seq; GSE91235 for A549 12hr-dex-treated GR ChIP-seq; GSE92032 for HeLa-S3 control ChIP-seq; GSE101280 for A549 REST control ChIP-seq; GSE101362 for A549 REST ChIP-seq.

### Processing of high-throughput sequence data

FASTQs files were aligned to the human genome reference assembly hg38 with bowtie2 (version 2.3.4.3; (Langmead and Salzberg 2012)), using the --sensitive option and requiring a MAPQ of at least 30. Fragments were discarded if they are aligned to gap, centromere, and telomere that are available in UCSC Gap and Centromere table browser (Hinrichs et al. 2006) and ENCODE blacklist regions (Amemiya et al. 2019). Alignment of paired-end datasets were further restricted to require properly paired alignments. All alignments were deduplicated using Picard (MarkDuplicates, version 2.14.0) prior to downstream analyses. Unnormalized and RPKM-normalized (--binSize 1) bigwig files were generated by bamCoverage subcommand in deeptools (version 3.0.1; (Ramirez et al. 2016)) using --extendReads. The reported average fragment length was used to extend reads when generating single-end bigwigs. Unnormalized and normalized bigwig files were used for CRADLE inputs files and for visualizing signals in genome browser tracks, respectively. A549 ATAC-seq FASTQs were processed as above but were aligned to hg19 and required a less stringent MAPQ score (≥5). Peaks were called for ChIP-seq datasets using MACS2 (Zhang et al. 2008) with a FDR threshold of 0.05. The coordinates of autosomal inhibitor-responsive regulatory elements previously reported in hg19 (Muerdter et al. 2018) were converted to hg38 with liftOver (Hinrichs et al. 2006).

### PER1 BAC library preparation and sequencing

Purified PER1 BAC (CH17-212C17; chr17:7981103-8192310) was harvested from *E. coli* using standard protocols. Following DNA shearing using the Covaris S2 system, the BAC DNAs were size-selected using solid phase reversible immobilization (SPRI) beads. STARR-seq insert libraries were prepared using the NEBNext DNA Library Prep Master Mix kit and 50 ng of template DNA. Adapted DNAs were enriched in triplicate reactions via 3, 6, 12, or 18 cycles of PCR using the NEB Q5 PCR kit. The resulting libraries were characterized on the Agilent Tape Station prior to 50-cycles of paired-end sequencing on the Illumina MiSeq platform. FASTQs were aligned as above.

### Data processing for bias covariates

To obtain DNA structure parameters for fragment-end bias, we estimated minor groove width (MGW) and propeller twist (ProT) for all 5mers (total 1,024 sequences) using DNAShape (Zhou et al. 2013). For Gibbs free energy parameters, we used the estimated Gibbs free energy for all dimers (Protozanova et al. 2004). For G-quadruplex structure parameters, we used bigwig files that reported stability of G-quadruplex structure in whole genome with accession code GSE63874 in GEO (Chambers et al. 2015). For mappability scores, we downloaded the human mappability score bigwig files for 36mer and 50mer (Derrien et al. 2012), using accession codes (ENCSR821KQV, ENCSR093EEM) in ENCODE (Consortium 2012; Davis et al. 2018). For those G-quadruplex structure bigwig and mappability bigwig files, the genomic coordinates were in hg19 assembly so we used liftOver tool (Hinrichs et al. 2006) to convert them to hg38.

### Measuring technical biases in STARR-seq libraries

To investigate fragment-end bias, we counted the frequency of 5-mers starting 2 bp upstream of the 5’ end of positive strands of fragments in STARR-seq input libraries (Johnson et al. 2018; Muerdter et al. 2018). We compared this the 5-mer frequency to that observed in the reference genome (hg38) excluding gap, centromere, and telomere that are available in UCSC Gap and Centromere table browser (Hinrichs et al. 2006) and ENCODE blacklist regions (Amemiya et al. 2019). To examine Gibbs free energy, G-quadruplex structure, and mappability bias, we binned human chromosome 1 into 500 bp windows using a 250 bp step. We estimated the amount of potential technical bias in a window by calculating the mean of per-base measure of those biases using previously reported values: Gibbs free energy value (Protozanova et al. 2004); the percent of mismatch for G-quadruplex structure bias (Chambers et al. 2015); mappability score (Derrien et al. 2012). This analysis was limited to the PER1 BAC when estimating Gibbs free energy bias in the PER1 BAC library.

### Correcting technical biases in STARR-seq

We used the technical bias covariates in a mixture general linearized model (GLM) with a Poisson distribution and log link to correct STARR-seq signals. An estimate of the 90th percentile of observed coverage in input libraries (*Isig*_*P*90_) was calculated using 1 kb bookended regions. To ensure the mixture GLM models effects across the range of observed signals, we trained the model using a structured sampling strategy to select bookended regions without replacement such that the final training set is approximately 10^6^ bases in length. We evenly partitioned the training set to fit regions with input signal above and below *Isig*_*P*90_. The set of regions below *Isig*_*P*90_, were further evenly partitioned into the following percentile bins of observed coverage: [0, 20); [20, 40); [40, 60); [60, 80); [80, 90). To ensure representation across the upper tail of the STARR-seq signal distribution, regions above *Isig*_*P*90_ were asymmetrically partitioned as follows: 62.5% of regions were evenly divided into the following percentile bins of observed coverage, [90, 92); [92, 94); [94, 96); [96, 98); [98, 99), while the remaining 37.5 % of regions were binned into the 99th percentile of coverage.

Single base positions with observed input signal (*Isig*_*pos*_) above and below *Isig*_*P*90_ were independently fit to the GLM. To predict the total bias effects at each single base position we used windows twice the length of the median fragment length (*L*) centered on the position of interest (Fig. S7). We assumed each position was covered by *L* number of hypothetical fragments of *L* bp length with each overlapping by a single base. We then multiplied the same bias covariates for all fragments in that window with each covariate to the power of a unique beta as below.

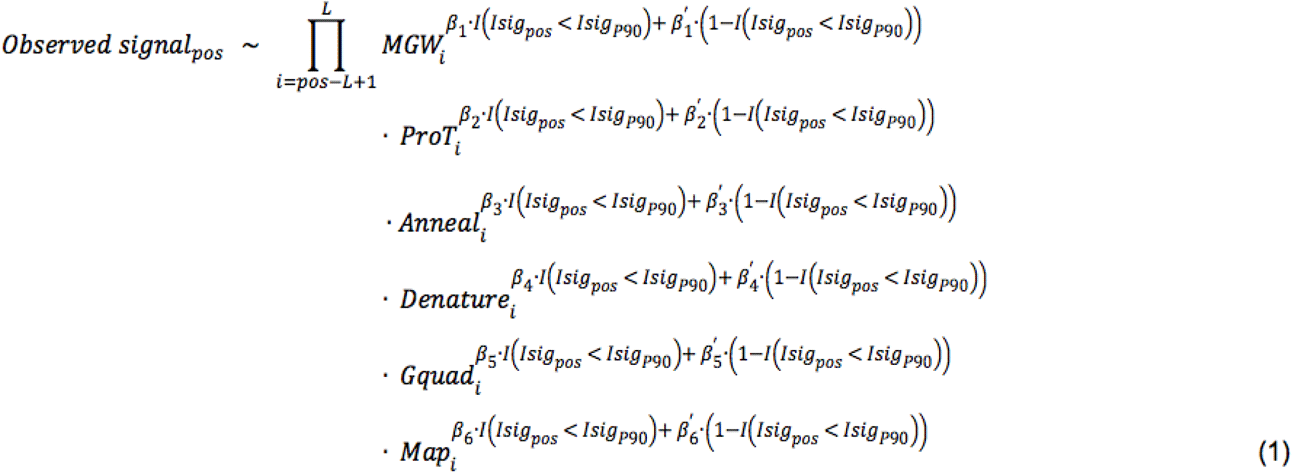

Each beta coefficient represents the relative effect of each bias predictor. Here, we assumed the set of betas is the same for all overlapping fragments. Then in log space, observed signal can be estimated with using the sums of bias covariates in the GLM as follows.

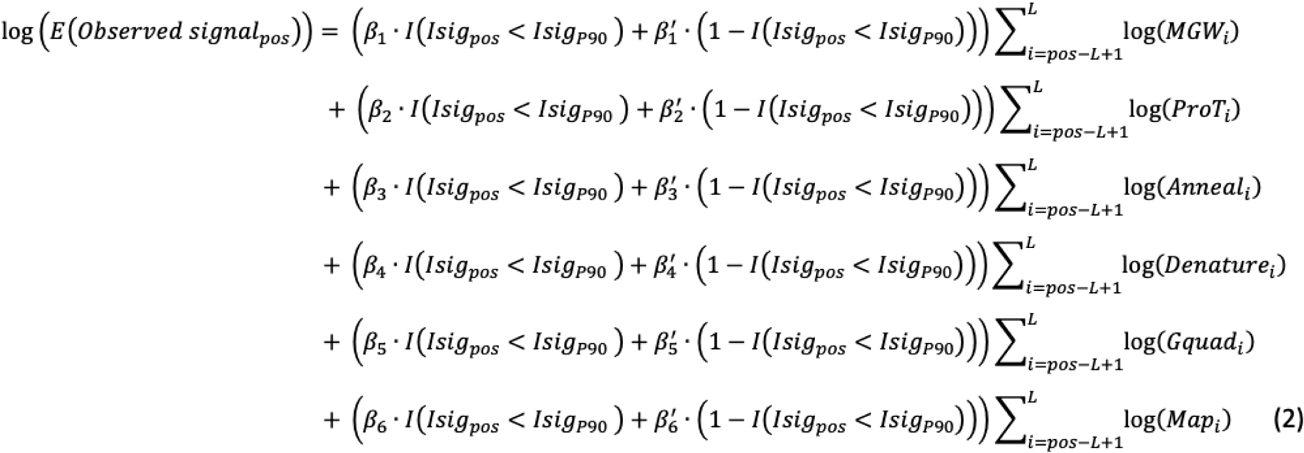

MGW and ProT values were calculated using DNAShape (Zhou et al. 2013) for the two 5mers starting from two bp external to both 5’ ends of each hypothetical fragment. *MGW*_*i*_ and *ProT*_*i*_. was obtained by multiplying those two MGW and ProT values, respectively.

We used the the nearest-neighbor model (SantaLucia 1998; Protozanova et al. 2004) to estimate the Gibbs free energy of each hypothetical fragment. To estimate the relative melting temperature (*T*_*m*_) of each fragment, we divided Gibbs free energy of a hypothetical fragment by the number of dimers in that hypothetical fragment and by the fixed entropy value (Protozanova et al. 2004). *T*_*m*_ values were normalized to range of [0, 1]. To model the non-linear dependency of anneal and denature efficiencies to *T*_*m*_, normalized *T*_*m*_ values in the *i*th fragment (*T*_*m,i*_) were mapped to two exponential functions as below.

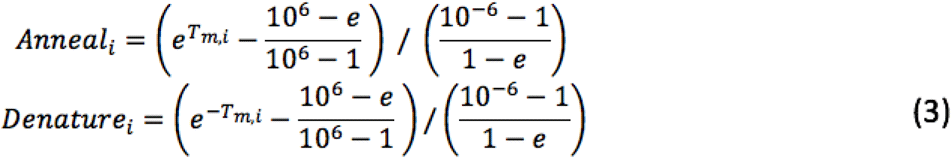

Those mapped values were used for *Anneal*_*i*_ and *Denature*_*i*_ in the GLM Model.

To obtain *Gquad*_*i*_ for each hypothetical fragment, we used the maximum of G-quadruplex structure stability value in that sequence (Chambers et al. 2015). To obtain *Map*_*i*_ for each hypothetical fragment, we used a kmer mappability score file (Derrien et al. 2012) where kmer is a sequencing length, and multiplied the mappability scores of both ends of a fragment.

After fitting the GLM, the bias predicted by the model at each base position was removed by subtracting the estimated bias effect from the observed signal. To avoid false positives, positions with fewer than ten observed overlapping fragments and with no signal in output libraries are not reported in the corrected signal file. The minimum number of observed overlapping fragments required for a position to be reported is parameterized (-mi) in the correctBias subcommand in CRADLE.

### Evaluating model fit

To determine how well the CRADLE GLM explained variance in observed signal, we calculated R^2^ with observed and predicted signals across chromosome 1 for each STARR-seq library. To calculate R^2^, we fitted the mixture GLM in CRADLE, and calculated the sum of squares (SSQ) by adding up squared residuals. Then, we calculated total SSQ, the sum of squared difference of observed signals and the mean signal. With using the SSQ and total SSQ, we calculated R^2^ with the following equation: *R*^2^ = 1 – (*SSQ*/*totalSSQ*).

### Evaluating the contribution of each covariate to model fit

To estimate the contribution of each bias type, we calculated semi-partial correlations for chromosome 1 using the mixture GLM. To assess each technical bias type, we excluded bias covariates that model corresponding bias type in fitting the GLM; for example, we excluded ‘anneal’ and ‘denature’ covariates when assessing Gibbs free energy bias impact. The R^2^ of these models were calculated as above and subtracted from the R^2^ of the full model.

### Calling regulatory elements with CRADLE

Genomic regions possessing regulatory activity were identified using a modified Benjamini method (Benjamini and Bogomolov 2014). We first binned the genome into windows (1.5x *L*) and determined the effect size of each window by subtracting the mean corrected signal in input libraries from the mean corrected signal in output libraries. Each window was classified to one of the three types, using the following standard:

Type(window_x_) = 1 if (effect size > 0 and |effect size| > 99th percentile of absolute effect sizes)

Type(window_x_) = -1 if (effect size < 0 and |effect size| > 99th percentile of absolute effect sizes)

Type(window_x_) = 0 else

The threshold of the 99th percentile of absolute effect sizes was chosen to classify windows because the majority of windows are not expected to encode regulatory activity. Contiguous windows of the same type, including Type 0, were merged to form regions for statistical testing. These regions were then binned with non-overlapping bins of which length is 1/6x *L.* In each bin, the input and output STARR-seq signals were compared using Welch’s t-test (Welch 1947) to account for potential differences in variance. Individual bin-level *P-*values from the same region were merged to a region-level *P-*value via the Simes’ method (RJ 1986). To increase our power to detect potential regulatory regions for final testing, regions with small overall variance were removed from further analysis, independently of the statistical test used (Bourgon et al. 2010). Specifically, we ranked regions according to their overall variance and then applied the overall variance filter that removed 0-90% of regions with low variance using 10% intervals. *P-*values for regions passing each threshold were subjected to the first BH procedure (Benjamini and Hochberg 1995) using a parameterized FDR value (-fdr). Then, we chose the threshold of the overall variance filter that returned the greatest number of selected regions from the first BH procedure. To identify bins that have regulatory activity, bin-level *P-*values from the Welch’s t-test in the selected regions were then subjected to the second BH procedure with new FDR adjusted by following equation: new FDR = (pre-determined FDR) x (the number of selected regions / total number of regions) Contiguous bins that encode regulatory activity with the same sign of effect sizes were merged in the final output and the minimum *P-*value was reported.

### Simulation of STARR-seq signals

To evaluate the performance of the CRADLE pipeline, we simulated STARR-seq signals that maintained the observed sequence biases and expected variance across replicates. STARR-seq signals were simulated using a negative binomial distribution and mean-variance relationships estimated independently for input and output libraries from previously published STARR-seq data (Johnson et al. 2018). Simulated input and output signal matrices, generated using 300 bp bookended windows along chromosome 1, were used to estimate mean-dispersion relationship in deseq2 (Love et al. 2014) prior to interpolation with the scipy.interpolate.interp1d command in python. To generate a set of pre-defined regulatory elements (N = 50,504), we randomly sampled ∼0.5% of total windows requiring that the selected windows to be in at least the 70th percentile of coverage in the published input libraries. For each pre-defined regulatory element we randomly assigned an absolute fold change [2, 3, 4] and regulatory activity type [activating, repressive]. Pre-defined sets of regulatory elements with a specific fold change and regulatory activity type were generated as above using the specified fold change and regulatory activity types as described in text.

Five simulated STARR-seq input and output signals were generated using a negative binomial distribution. The mean parameters used to generate the simulated input and output signals were determined by calculating the mean window counts using the published input libraries (Johnson et al. 2018). The variance parameters were determined using either the input or output interpolation analyses described above. The mean parameters used to generate the simulated output signals were adjusted for pre-defined regulatory elements windows by multiplying or dividing the mean signal by the pre-determined fold change and determining the corresponding variance parameter.

### Detecting regulatory elements in simulated data

To evaluate the effect of correcting STARR-seq signals on identifying regulatory elements, we used CRADLE to call regulatory elements before and after correcting biases in the simulated datasets. Due to the normality assumption in Welch’s t-test, we modified the CRADLE approach described above to call regulatory activity in uncorrected simulated signals. In place of the Welch’s t-test, we used two alternative statistical approaches to compare uncorrected simulated input and output signals. First, we used a Poisson GLM as follows:

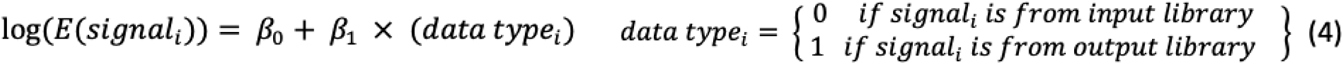

We then performed the Wald test for *β*_1_ with f distribution. Second, we followed a similar approach as used by MACS2 (Zhang et al. 2008). We used the mean input bin signal as the mean parameter in a Poisson distribution to calculate a *P-*value for the mean output bin signal. We called regulatory activity in corrected simulated signals as described above.

### Motif enrichment analysis

Motif enrichment analysis was performed using the findMotifsGenome subcommand in the Homer (Heinz et al. 2010) 4.10.1 software suite using the following parameters: -size given -mis 3 -mset vertebrates.

### TF occupancy in CRADLE regulatory elements

To determine whether regulatory elements called by CRADLE are bound by TFs, we used the CRADLE pipeline described above to detect A549 steady-state activating and repressive regulatory elements from previously published study (Johnson et al. 2018). We used the findMotifsGenome subcommand in the Homer suite (version 4.10.1; (Heinz et al. 2010)) with the following parameters, -size given -mis 0 -mset vertebrates -find, to detect REST motifs in each repressive element and FOSL2, JUNB, and GABPA motifs in activating elements. For each element that encoded a specified motif, we intersected those elements with ENCODE ChIP-seq peaks for the corresponding TF in A549 cells (Consortium 2012; Davis et al. 2018).

### Validation of REST occupied repressive regulatory elements

Regions tested by Jayavelu et al. in K562 cells (N = 7,440; (Doni Jayavelu et al. 2020)) were intersected with repressive regulatory elements identified by CRADLE in A549 cells that also contained a REST motif and were bound by REST in the same cell line (N = 240) (Consortium 2012). Reported fold change values in K562 cells were compared for the intersection set except the two elements without coverage (N = 28), regions predicted by Jayavelu et al. to be repressive elements (N = 3,001), and control regions (N = 40).

Whole genome survey of regulatory elements (SuRE) signals from HepG2 and K562 cells (van Arensbergen et al. 2019) were compared in specific sets of regulatory elements identified by CRADLE in A549 cells. These regulatory elements included activating regulatory elements that contained either a FOSL2, GABPA, or JUNB motif and were bound by the corresponding TF in A549 (Consortium 2012) or repressive elements that likewise contained a REST motif and were bound by REST (N = 240; (Consortium 2012)). A549 regulatory elements that contained a SNP in the genomes assayed in the SuRE study were excluded on a per genome basis. The minimum and maximum number of SNP-filtered elements compared for each TF are as follows: FOSL2 N=650-651; GABPA N=401-402; JUNB N=723; REST N=102.

We randomly generated a set of regions (N = 240) of fixed length (430 bp) controlling for accessibility (Consortium 2012) and dinucleotide composition. The fixed length was set to the median length of the compared repressive elements. In generating random regions, we excluded regions that overlapped gaps, centromeres, and telomeres that are available in UCSC Gap and Centromere table browser (Hinrichs et al. 2006) and ENCODE blacklist regions (Amemiya et al. 2019), or the following features defined by chromHMM (Ernst and Kellis 2017) in K562 and HepG2 cells: promoters, promoter flanking regions, enhancers, CTCF enriched sites, and repressed regions. After applying the SNP-filter described above, we obtained 94 random regions.

## Data access

The PER1 BAC datasets generated during the current study are available in GEO with accession number GSE149914.

CRADLE is implemented in Python and freely downloadable either from github (https://github.com/Young-Sook/CRADLE) or pip (pip install cradle). Instructions for installing and running CRADLE are available on the CRADLE github page.

## Acknowledgments

We thank Greg Crawford and David MacAlpine for their helpful comments and advice in developing this work.

## Author Contributions

Conceptualization and project administration was by Y.K. and T.E.R.; data curation, formal analysis, investigation, validation, and visualization was by Y.K.; supervision and funding acquisition was by T.E.R.; methodology was by Y.K., T.E.R, G.D.J., J.S., W.H.M, A.O., and A.S.A.; software development was by Y.K. and A.B.; writing of the original draft was by Y.K. and T.E.R; and review and editing was by all of the authors.

## Disclosure declaration

The authors declare that they have no competing interests.

## Notes

### Competing Interest Statement

The authors have declared no competing interest.

