## Supplemental_Fig_S1 for "Detecting regulatory elements in high-throughput reporter assays"

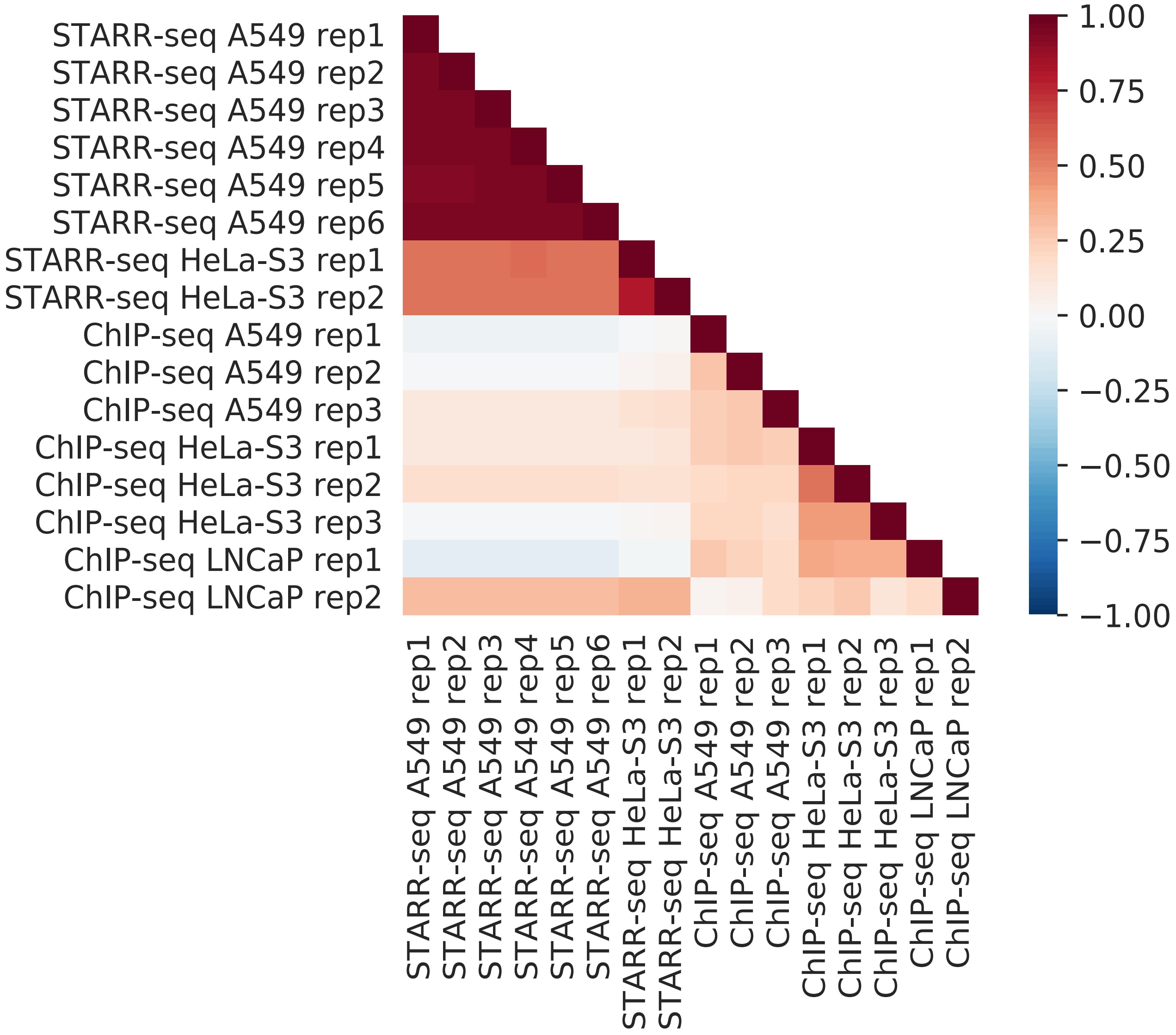


**Figure S1** | Pearson correlations of 1 bp signals in STARR-seq input [5, 6] and ChIP-seq control [7] libraries along chromosome 1.
