## Supplemental_Fig_S2 for "Detecting regulatory elements in high-throughput reporter assays"

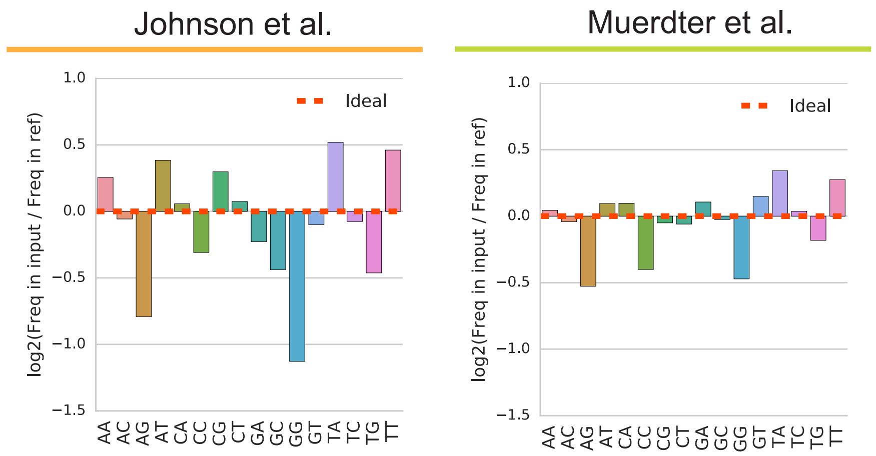


**Figure S2** | DNA structure bias in the terminal positions of STARR-seq fragments. The frequencies of observed dimers starting from one bp external to the 5' ends of fragments was compared to that in reference autosomes (hg38) excluding excluding gap, centromere, and telomere that are available in UCSC Gap and Centromere table browser [34] and ENCODE blacklist regions [35].
