## Supplemental_Fig_S3 for "Detecting regulatory elements in high-throughput reporter assays"

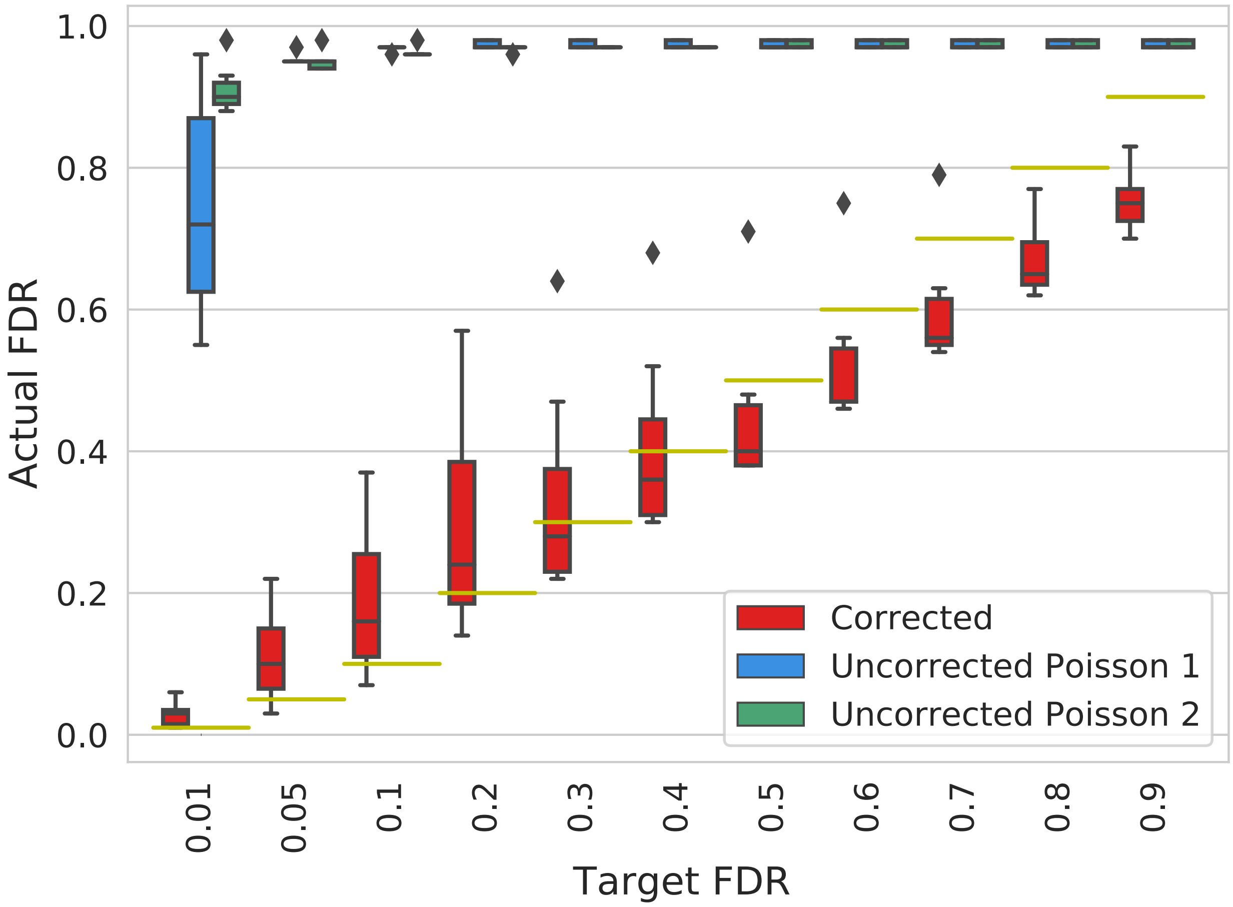


**Figure S3 |** Determination of actual FDR for regulatory elements detected by CRADLE using simulated STARR-seq data. The relationship between parameterized target FDR values and actual FDR values calculated using simulated corrected (red) and uncorrected STARR-seq signals. To detect regulatory elements with uncorrected signals, two statistical approaches were used (see methods; ‘Uncorrected Poisson1’, blue; ‘Uncorrected Poisson2’, green). The yellow line indicates target FDR. Whiskers extend 1.5 times the interquartile range. Center lines show the medians.
