## Supplemental_Fig_S4 for "Detecting regulatory elements in high-throughput reporter assays"

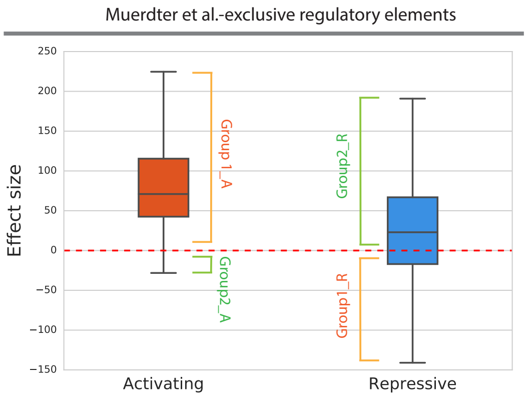


Moitfs enriched in Group1_A compared to Group2_A

| Rank | Motif | Name | P value |
|---|---|---|---|
| 1 | GATGAGTCATCC | FOSL2 | $10^{-404}$ |
| 2 | GCAACAGGTG | SCRT1 | $10^{-350}$ |
| 3 | GGAAATTCCC | NFkB-p64-Rel | $10^{-333}$ |

Moitfs enriched in Group2_A compared to Group1_A

| Rank | Motif | Name | P value |
|---|---|---|---|
| 1 | GAAAGTGAAAGT | IRF2 | $10^{-2}$ |
| 2 | TCATCAATCA | PDX1 | $10^{-2}$ |

Moitfs enriched in Group1_R compared to Group2_R

| Rank | Motif | Name | P value |
|---|---|---|---|
| 1 | AGTTTCAGTTTC | IRF3 | $10^{-59}$ |
| 2 | GAAAGTGAAAGT | IRF2 | $10^{-57}$ |
| 3 | AGTTTCAGTTTC | ISRE | $10^{-53}$ |

Moitfs enriched in Group2_R compared to Group1_R

| Rank | Motif | Name | P value |
|---|---|---|---|
| 1 | AGGGGATTTCCC | NFkB-p65 | $10^{-27}$ |
| 2 | GATGACTCATCC | Jun-AP1 | $10^{-17}$ |
| 3 | AAATTGCTGAGTCAT | BACH1 | $10^{-15}$ |

**Figure S4 |** CRADLE more accurately estimates regulatory element effect sizes. CRADLE effect sizes were plotted for regulatory elements exclusively called by Muerdter et al. [6]. Activating and repressive regulatory elements were subsetted according to the sign of their effect size into Group1 and Group2. Group1 subsets includes activating regulatory elements with a positive effect size (Group1_A) and repressive regulatory elements with a negative effect size (Group1_R). In contrast, Group 2 subsets includes activating regulatory elements with a negative effect size (Group2_A) and repressive regulatory elements with a positive effect size (Group2_R). Motif enrichment analysis was performed for each group relative to its partner.
