## Supplemental_Fig_S5 for "Detecting regulatory elements in high-throughput reporter assays"

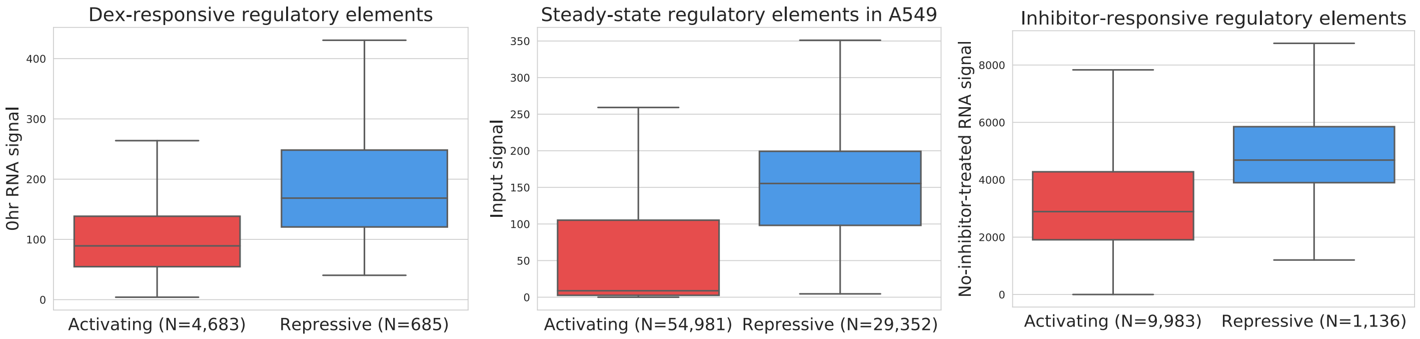


**Figure S5 |** Distribution of control library signals for regulatory elements detected by CRADLE in STARR-seq studies [5, 6]. Whiskers extend 1.5 times the interquartile range. Center lines show the medians.
