## Supplemental_Fig_S6 for "Detecting regulatory elements in high-throughput reporter assays"

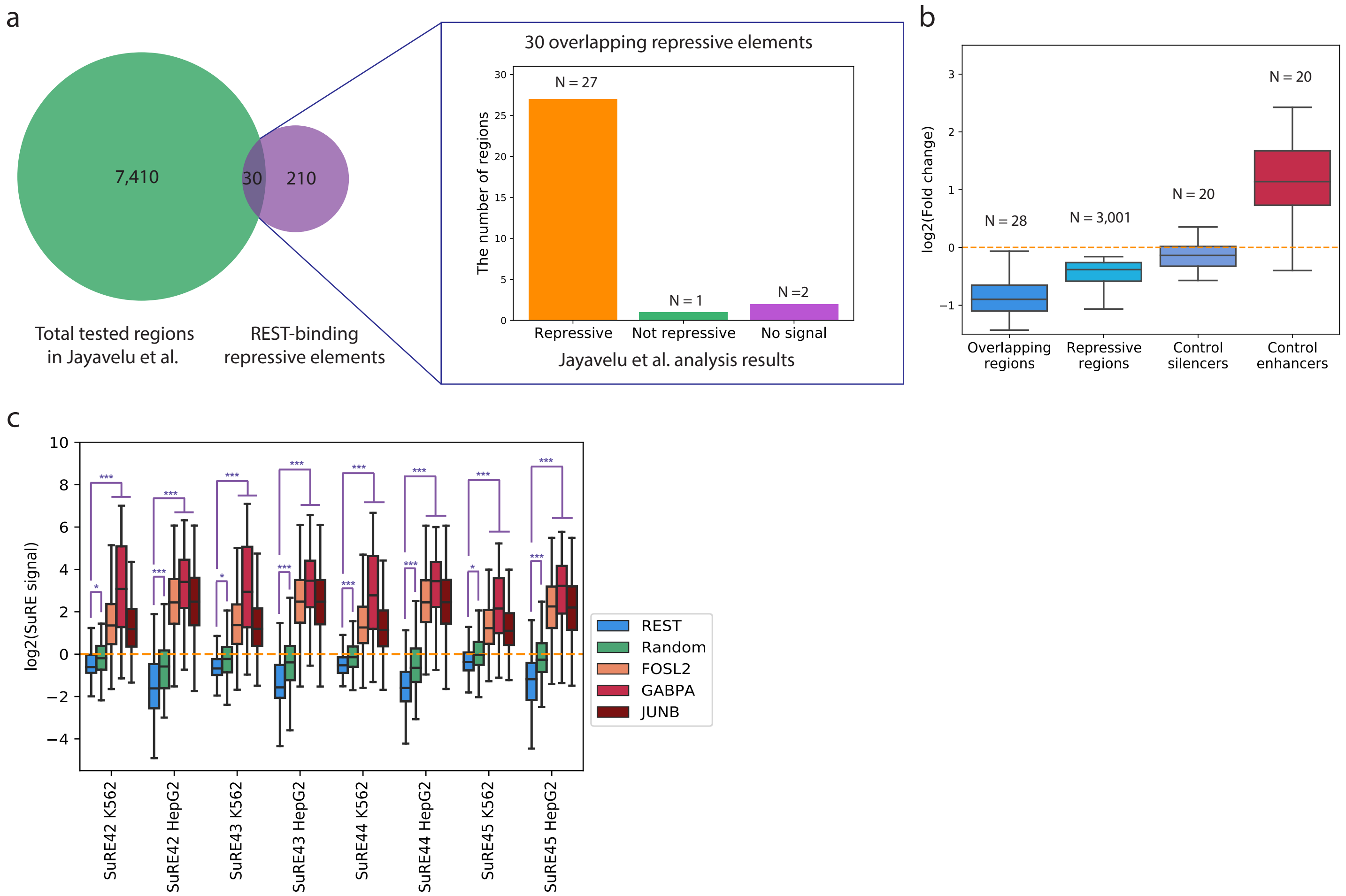


**Figure S6** | Validation of REST-binding A549 steady-state repressive regulatory elements identified by CRADLE. a, Venn diagram showing the intersection of tested regions in Jayavelu et al. [30] and steady-state REST-binding repressive elements in A549 cells. Among the 30 elements in the intersection, 27 elements were previously reported to be repressive. b, The distribution of previously-reported fold changes for the elements in the intersection as well as repressive and control regions in the prior study. The two elements without coverage in the intersection were not included. c, Whole genome survey of regulatory elements (SuRE) signals from HepG2 and K562 cells [31] were compared in subsets of regulatory elements identified by CRADLE in A549 cells. These regulatory elements included activating regulatory elements that contained either a FOSL2, GABPA, or JUNB motif and were bound by the corresponding TF in A549 [7], repressive elements that likewise contained a REST motif and were bound by REST [7], or a set of randomly generated regions. Whiskers extend 1.5 times the interquartile range. Center lines show the medians. Wilcoxon rank-sum test was used for statistical testing. P-values from: ***, P-value < 0.0001; *, 0.001 < P-value < 0.01.
