## Supplemental_Fig_S7 for "Detecting regulatory elements in high-throughput reporter assays"

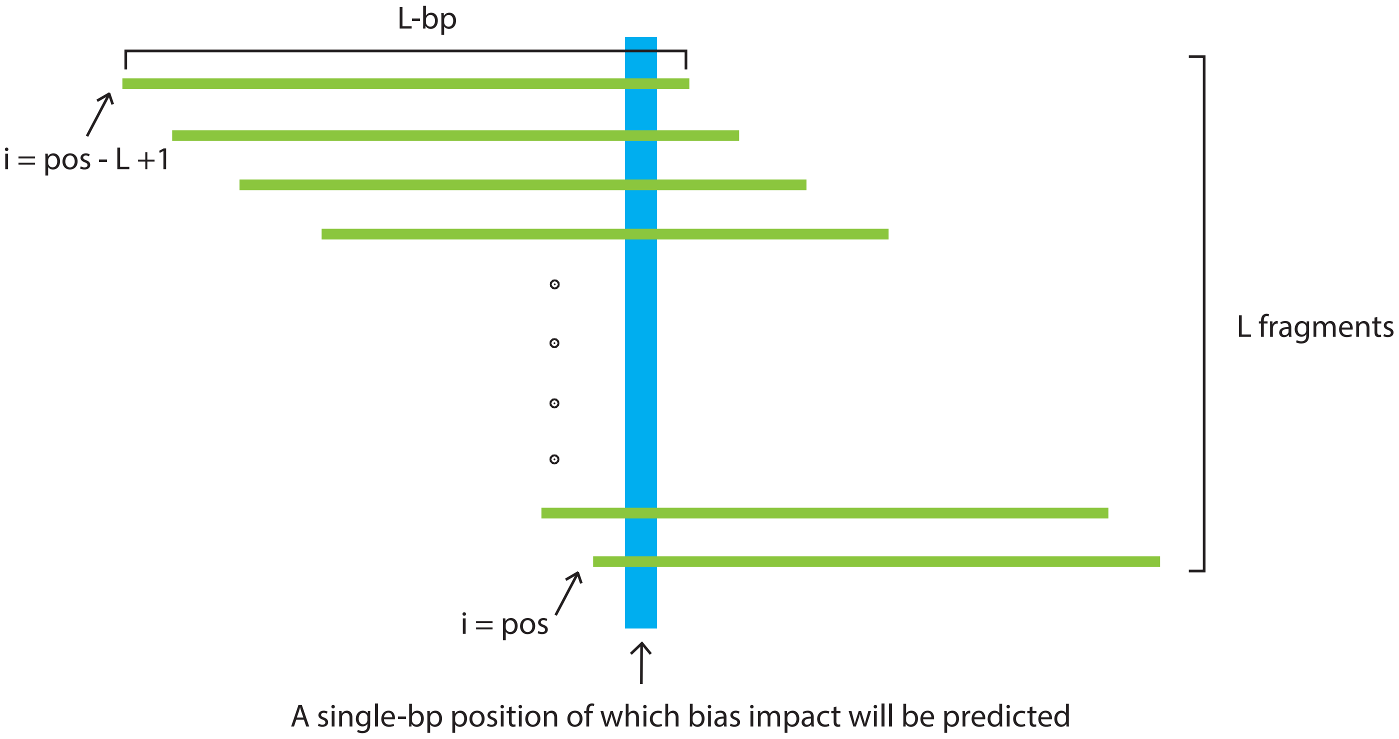


**Figure S7** | Approach used by CRADLE to calculate bias covariates. To estimate bias effects for each position (blue), we used a window centered on that position that was twice the median fragment length, L. We assume L number of fragments (green) in a window and that each fragment is L-bp in length.
